# Structured RhoGEF recruitment drives myosin II organization on large exocytotic vesicles

**DOI:** 10.1101/2023.09.06.556511

**Authors:** Kumari Kamalesh, Dagan Segal, Ori Avinoam, Eyal D. Schejter, Ben-Zion Shilo

## Abstract

The Rho family of GTPases plays a crucial role in cellular mechanics, by regulating actomyosin contractility through the parallel induction of actin and myosin assembly and function. Using exocytosis of large vesicles in the *Drosophila* larval salivary gland as a model, we followed the spatiotemporal regulation of Rho1, that in turn creates distinct organization patterns of actin and myosin. After vesicle fusion, low levels of activated Rho1 diffuse to the vesicle membrane and drive actin nucleation in an uneven, spread-out pattern. Subsequently, the Rho1 activator RhoGEF2 distributes as an irregular meshwork on the vesicle membrane, activating Rho1 in a corresponding punctate pattern and driving local myosin II recruitment, resulting in vesicle constriction. Vesicle membrane buckling and subsequent crumpling occur at local sites of high myosin II concentrations. These findings indicate that distinct thresholds for activated Rho1 create a biphasic mode of actomyosin assembly, inducing anisotropic membrane crumpling during exocrine secretion.

## Introduction

The Rho subfamily of small GTPases is a focal point for controlling cytoskeletal architecture and cell behavior in eukaryotic cells ^(Hodge & Ridley, 2016; Jaffe & Hall, 2005)^. Rho proteins are loosely anchored to the cell membrane via C-terminal lipid modifications. Both their upstream regulators and downstream effectors are cytoplasmic proteins recruited to the membrane by their transient association with Rho. Rho proteins shuttle between the active, GTP-bound, and inactive GDP-bound states. RhoGEF activators and RhoGAP inhibitors orchestrate the transition between these states. The spatial and temporal dynamics of these two classes of regulators, which vary according to the cell and tissue requirements, determine the resulting pattern of Rho activation(Bos *et al*, 2007; Denk-Lobnig & Martin, 2019). Activated Rho GTPases recruit downstream target proteins to the membrane, triggering a cascade of events that reorganize the cytoskeleton, modify cell adhesion and motility, and activate or repress gene expression (Spiering & Hodgson, 2011).

Upon activation, Rho-GTP catalyzes force generation through the actomyosin meshwork. This is accomplished by recruiting and activating Formin family proteins that execute the generation of linear actin microfilaments, and myosin II motors that link to anti-parallel actin cables for force generation. Formins are present in the cytoplasm in an inactive state due to the intra-molecular association between the N- and C-termini of the protein. Upon binding to membrane-associated Rho-GTP, Formins transition from a closed to an open state and become active. Stable association with the membrane is mediated by binding to the phospholipid PtdIns (4,5)P_2_ (PIP2) via a basic domain in the Formin protein (Rousso *et al*, 2013). The combined association with Rho-GTP and PIP2, termed “coincidence detection”, leads to a stable membrane interaction of active Formin and the nucleation of linear actin microfilaments.

The recruitment and binding of non-muscle myosin II to the actin cables is driven by Rho-associated protein kinase (Rock). Like Formins, Rock is a cytoplasmic protein maintained in an inactive state due to the intra-molecular association between its N- and C-termini. Following the initial recruitment by active Rho-GTP, a PH phospholipid-binding domain in Rock provides tighter association with the membrane (Amano *et al*, 2000). Rock phosphorylates the regulatory light chains of the myosin II hetero oligomer, allowing the complex to bind the actin microfilaments. Thus, Rho-GTP is the common trigger for both actin polymerization and myosin II activation. However, how the same molecule temporally and spatially coordinates these two distinct events remains unclear.

Actomyosin plays a central role during exocrine secretion processes that employ large secretory vesicles (Nightingale *et al*, 2012). A prominent example are the *Drosophila* larval salivary glands (LSGs), which utilize large secretory vesicles (SVs), several microns in diameter, and display an orchestrated recruitment of actomyosin, which is required in order to expel their content into the lumen of the gland. Salivary gland SVs release large amounts of adhesive mucinous glycoproteins (“Glue”), which the fly pupa uses to attach to a solid surface. Secretion initiates shortly before pupation in response to a hormonal (Ecdysone) signal (Biyasheva *et al*, 2001), proceeds for approximately 2 hours, and involves dozens of individual secretory vesicles within each cell. Exocytosis of each vesicle takes 1-3 minutes (Rousso *et al*, 2016; Tran *et al*, 2015). Interestingly, actomyosin contractility mediates folding and “crumpling” of the vesicle membrane, which drives a unique mode of exocytosis, wherein content release occurs while the vesicle membrane remains segregated from the apical cell membrane (Kamalesh *et al*, 2021). Activation of Rho1, the primary *Drosophila* Rho family homolog, triggers both arms of the actomyosin network in this setting-actin polymerization and myosin II recruitment and activation (Rousso *et al*., 2016). While previous work identified RhoGAP71E as a RhoGAP protein recruited to the secretory vesicles to drive Rho1 inactivation and actin depolymerization (Segal *et al*, 2018), the corresponding RhoGEF proteins mediating Rho1 activation have not been identified. Moreover, the precise mechanism of actomyosin organization and force generation on the SVs has not been established.

In the current study, we examine the spatiotemporal dynamics of the Rho network during exocytosis of large secretory vesicles, by exploring the activation of Rho1 by RhoGEF proteins and its implication on actomyosin contractility. We show that upon vesicle fusion with the apical cell membrane, low levels of activated Rho1 appear on the vesicle membrane. These levels are sufficient to recruit the Formin Diaphanous (Dia), generating a fairly uniform F-actin coat around the vesicle (Segal *et al*., 2018). However, this initial Rho-GTP pulse does not recruit sufficient levels of myosin II to mediate vesicle constriction. Subsequent recruitment of RhoGEF2 in an irregular structured pattern on the F-actin coat, generates a corresponding activation pattern of Rho1, driving local myosin II recruitment and subsequent constriction. Vesicle membrane buckling occurs at sites of high myosin II concentration, leading to the crumpled shape of the vesicle membrane during content release. Our finding points to a paradigm of different thresholds for recruitment of Dia and Rock by activated Rho1, enabling a biphasic mode of actomyosin recruitment that drives anisotropic membrane crumpling during large secretory vesicle exocytosis.

## Results

### Distinct patterns of recruitment and organization of F-actin and myosin II on secretory vesicles

In the *Drosophila* LSGs, the fusion of large secretory vesicles with the apical cell membrane initiates an orchestrated process of actomyosin recruitment to the vesicle membrane that expels its content (Fig. 1 A, B). To characterize actomyosin coat dynamics and organization by Rho1 signaling, we monitored the F-actin probe LifeAct-Ruby and the myosin II light-chain reporter Sqh-GFP, using Airyscan confocal live-imaging (Huff, 2015) of ex-*vivo* cultured LSGs. F-actin and myosin II appeared almost simultaneously on the vesicles approximately 40s after fusion, but displayed distinct spatial assembly patterns (Fig. 1 C, D). The actin coat adopted a fairly uniform, spread-out distribution, while myosin II appeared punctate (Fig. 1 C, D).

**Figure 1.**
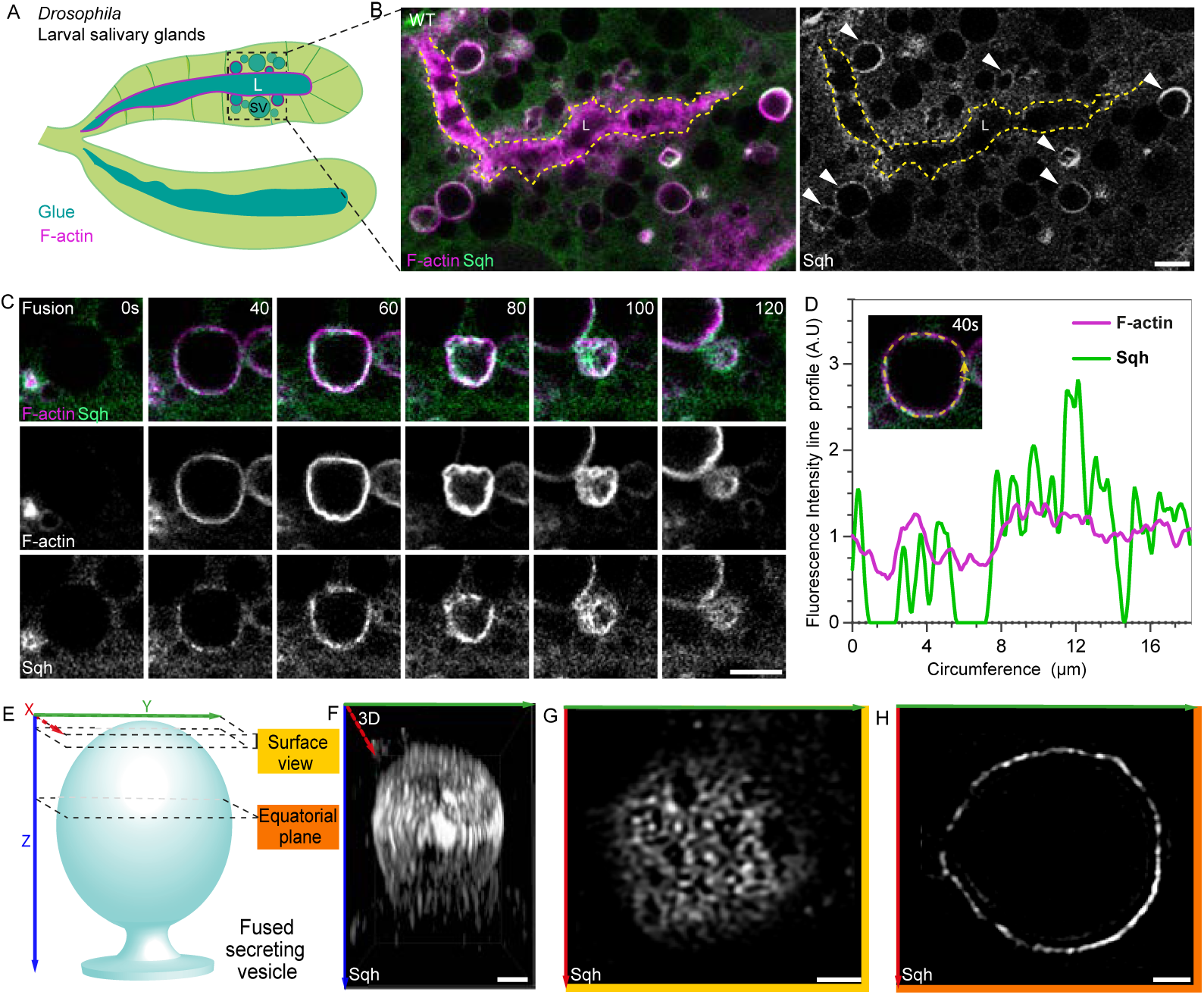
Distinct patterns of actin polymerization and myosin II recruitment on secretory vesicles. A. Schematic of *Drosophila* third instar larval salivary glands (LSGs). The gland cells (light green) are loaded with large secretory vesicles (SVs) that are filled with glue proteins (dark green). These SVs fuse with the apical membrane and release their content to the lumen (L). Vesicle fusion is followed by the assembly of an F-actin coat (magenta), and an orchestrated cycle of actomyosin organization leads to constriction of the vesicle and content release. B. Imaging live *ex vivo* cultured LSGs using Airyscan microscopy. SVs fuse with the apical membrane (dashed yellow line), and initiate assembly of F-actin (magenta, visualized using LifeAct-Ruby) and recruitment of myosin II (green or gray, visualized using Sqh-GFP, a GFP-tagged variant of Spaghetti squash (Sqh), the *Drosophila* myosin II regulatory light chain homolog). C. Airyscan time series of actomyosin on a representative vesicle following fusion with the apical membrane, using LifeAct-Ruby (magenta-top panel or gray-middle panel) and Sqh-GFP (green-top panel or gray-bottom panel). D. F-actin and myosin II fluorescence intensity line profiles along the circumference of the vesicle from panel C at 40 seconds (s) post-fusion, normalized to the mean intensity. Arrowhead in the inset marks the beginning of the line profile. F-actin showed a relatively uniform spread-out distribution, while myosin II was recruited in distinct puncta. E-H. 3D organization of myosin II as observed by Airyscan super-resolution imaging of fixed LSGs, following anti-GFP immunostaining for Sqh-GFP. (E) Shows a scheme of the position of imaging planes of the vesicle used for representation or analysis. The surface view is the 3D rendering of myosin II organization on any surface of the vesicle, using a Z-projection from the first 6 slices (details in methods section). Myosin II was visible as a discrete heterogenous meshwork on the vesicle surface. Individual planes of the Z-series of the image in (G) are shown in Figure S1. The equatorial plane is a slice at the middle of the vesicle (image shown in (H)). (F) Shows the 3D distribution of Sqh-GFP as seen from a side view of a fused vesicle represented by the scheme in (E). Scale bars: 5 µm (B-C), 1 µm (F-H)

To obtain a comprehensive 3D view of actomyosin distribution on the entire vesicle, we examined fixed LSGs stained for Sqh-GFP and Phalloidin, using Airyscan confocal super-resolution microscopy and deconvolution (Fig. 1 E-H). Examining the vesicles at their equatorial planes showed myosin II localization as a punctate pattern, consistent with the live imaging (Fig. 1 C-E and H). 3D rendering from the vesicle surface revealed a more complete picture - a structured meshwork-like organization of myosin II on the vesicle (Fig. 1 E-G and Fig. S1 A, B, C i, iii), alongside a more spread-out arrangement of F-actin (Fig. S1 C i, ii). These observations suggest that while activated Rho1 is the common trigger, additional factors contribute to the distinct spatial patterning of F-actin and myosin II on SVs.

### RhoGEF2 is essential for upregulating levels of activated Rho1 on secreting vesicles

To examine the basis for the different recruitment patterns of F-actin and myosin II to SVs, we searched for the RhoGEF protein(s) responsible for activating Rho1 in this setting. One prominent candidate was RhoGEF2, known for its role in activating Rho1 in a variety of *Drosophila* tissues (Fox & Peifer, 2007; Grosshans *et al*, 2005; Hacker & Perrimon, 1998; Mulinari *et al*, 2008; Padash Barmchi *et al*, 2005). To test the role of RhoGEF2 in this context, we monitored the distribution of the active Rho sensor AniRBD::GFP (Munjal *et al*, 2015) and F-actin following RNAi-mediated knockdown of RhoGEF2 in LSGs (Fig. 2 A, B). In wild-type (WT) SVs, the Rho sensor and F-actin are both typically observed on the vesicles within 40s after fusion and persist until exocytosis completes (Fig. 2 C). The Rho sensor initially appeared faint and spread-out, but subsequently intensified and became punctate (Fig. 2 C). Notably, the Rho sensor was occasionally observed before F-actin, consistent with the notion that Rho activation precedes actin recruitment.

**Figure 2.**
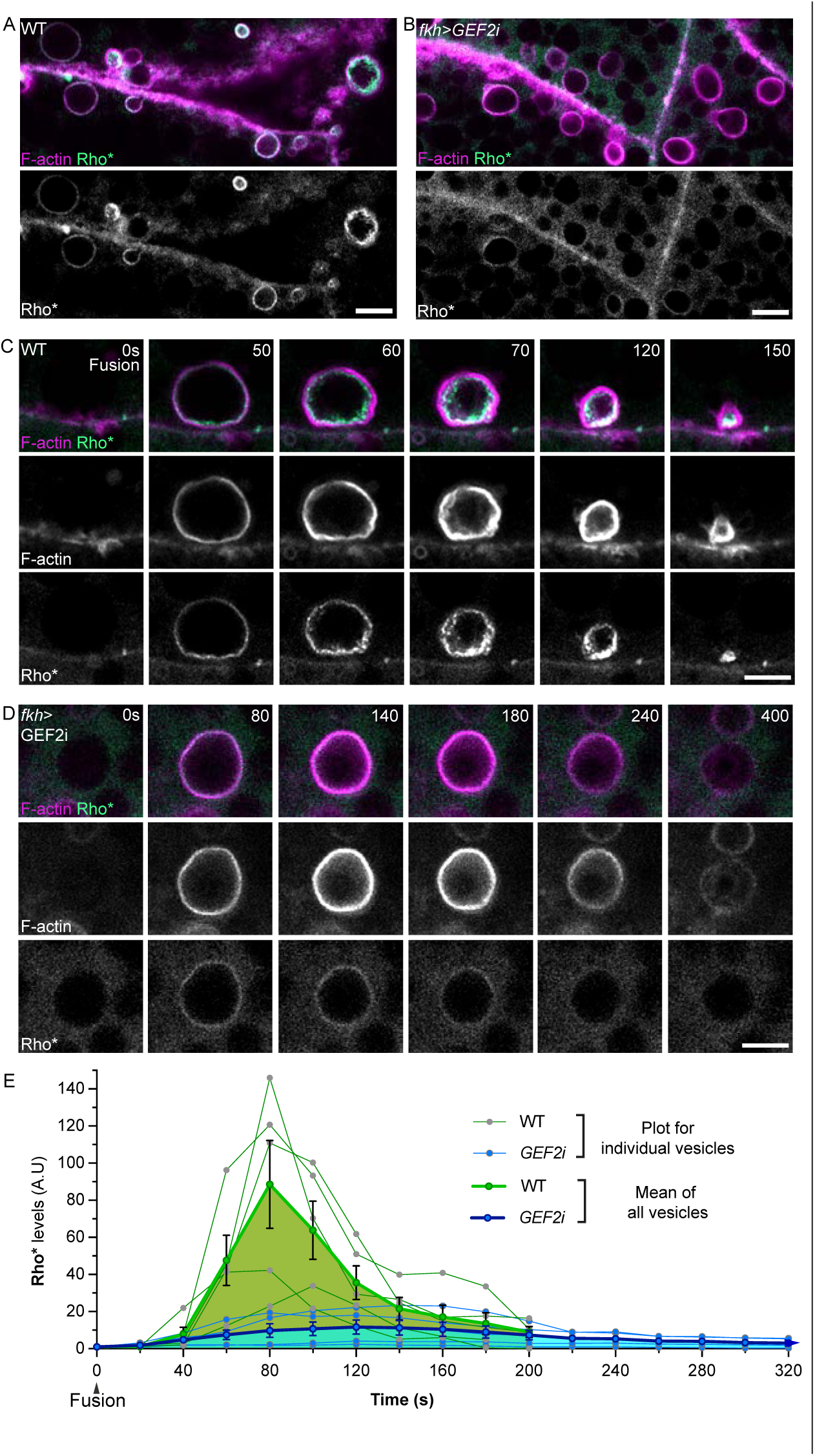
RhoGEF2 is essential for upregulating levels of activated Rho1 on secreting vesicles. A-B. Live imaging of *ex vivo* cultured WT and *RhoGEF2* knockdown (*fkh-Gal4>UAS-RhoGEF2* RNAi) LSGs. Abnormally low levels of the active Rho sensor Ani-RBD-GFP (Rho*, green-top panel or gray-bottom panel) were observed on F-actin coated SVs (LifeAct-Ruby, magenta) in LSGs expressing *RhoGEF2* RNAi. C. Time series of activated Rho1 and F-actin dynamics. Appearance of F-actin and active Rho sensor were observed earlier than 50 seconds post-fusion, and persisted throughout the constriction of the vesicle. The activated Rho signal dynamically increased and appeared highly punctate during the course of secretion. D. Time series of a *RhoGEF2* knockdown (*fkh-Gal4>UAS-RhoGEF2* RNAi) vesicle. In these glands, F-actin assembly on SVs was normal. However, the levels of active Rho sensor were low, and the vesicles failed to constrict or showed a significant delay in constriction. Secretion was stalled in 75±8% of the vesicles (n=75 vesicles from 3 LSGs). The typical F-actin disassembly cycle also appeared aberrant in this stalled vesicle. E. Plot of total active Rho levels on fused SVs over time. 0s marks vesicle fusion. Rho levels were normalized to the first appearance of a signal (typically at 30s). Images were acquired at 5s intervals. Plots of individual SVs are shown for WT and *RhoGEF2* RNAi (green and blue thin lines, respectively). The mean intensity profile (WT and *RhoGEF2* RNAi n=5 vesicles, from 3 LSGs, error bar SEM) is shown by a green and blue fill, respectively. Active Rho levels were dramatically reduced in *RhoGEF2* RNAi vesicles. Scale bars: 5 µm (A-D).

In RhoGEF2 RNAi-expressing LSGs, the SVs exhibited prominent actin recruitment comparable to WT. However, the Rho sensor appeared weak and diffuse (Fig. 2 D and Fig. S2 A). Quantitative analysis of multiple vesicles highlights the significantly reduced levels of the recruited Rho sensor on SVs in RhoGEF2 RNAi-expressing glands, as compared to WT LSGs (Fig. 2 E). Consistent with the reduced levels of active Rho, most of the fused vesicles stalled and failed to expel their content (75±8%, n=75 SVs, N=3 SGs; Fig. 2 D and Fig. S2 A). Furthermore, of the stalled vesicles, 78% also exhibited loss of the typical F-actin assembly and disassembly cycles observed in WT SVs that were halted by Rock inhibition. Our previous work showed that RhoGAP71E activity on stalled vesicles is responsible for this cycling of F-actin levels (Segal *et al*., 2018). The disassembly of F-actin, mediated by RhoGAP71E, was thus also affected by knockdown of RhoGEF2 activity. These effects were specific to RhoGEF2 RNAi, as expression of an RNAi construct targeting another highly expressed RhoGEF in LSGs, GEFmeso, displayed normal F-actin and Rho1 sensor recruitment, similar to WT (Fig. S2 B).

Based on these observations, we conclude that RhoGEF2 is essential for enhancing the levels of active Rho1 on SVs, which in turn impacts vesicular constriction. Furthermore, we find that F-actin recruitment to SVs is independent of the RhoGEF2-driven recruitment of Rho1 to SVs.

### RhoGEF2 is essential for myosin II recruitment and organization on the vesicles

The stalling of vesicular constriction, observed in the *RhoGEF2*-RNAi LSGs, prompted us to examine the recruitment and distribution of myosin II under these conditions. To this end, we monitored the F-actin probe LifeAct-Ruby and the myosin II light-chain reporter Sqh-GFP in *RhoGEF2* RNAi-expressing LSGs. We observed that myosin II recruitment and distribution were aberrant. The levels of recruited myosin II were significantly lower, and the pattern appeared more diffuse on the RhoGEF2 RNAi SVs (Fig. 3 A, B and Fig. S3 A; compare to WT in Fig. 1 C). Quantification of several vesicles showed the reduced myosin II levels in RhoGEF2 RNAi compared to WT LSGs (Fig. 3 C). Examination of the overall 3D organization of Sqh-GFP in fixed and immunostained LSGs revealed apparent differences in the myosin II pattern in RhoGEF2-RNAi SVs compared to WT (Fig. 3 F vs E). A 3D rendering of the myosin II organization from the surface of the vesicle revealed a diffuse distribution on RhoGEF2-RNAi SVs, unlike the structured organization observed in WT (Fig. 3 F i, iii vs. Fig. E i). Nevertheless, some intermittent gaps could be visualized in most vesicles. Consistently, Sqh-GFP also appeared uniform and diffuse at the equatorial plane of the *RhoGEF2* knockdown SVs (Fig. 3 F ii, iv vs Fig. 3 E ii). Expression of an RNAi construct targeting *GEFmeso* did not affect myosin II recruitment levels or the punctate distribution, which were similar to WT (Fig. 3 C and Fig S3 B).

**Figure 3.**
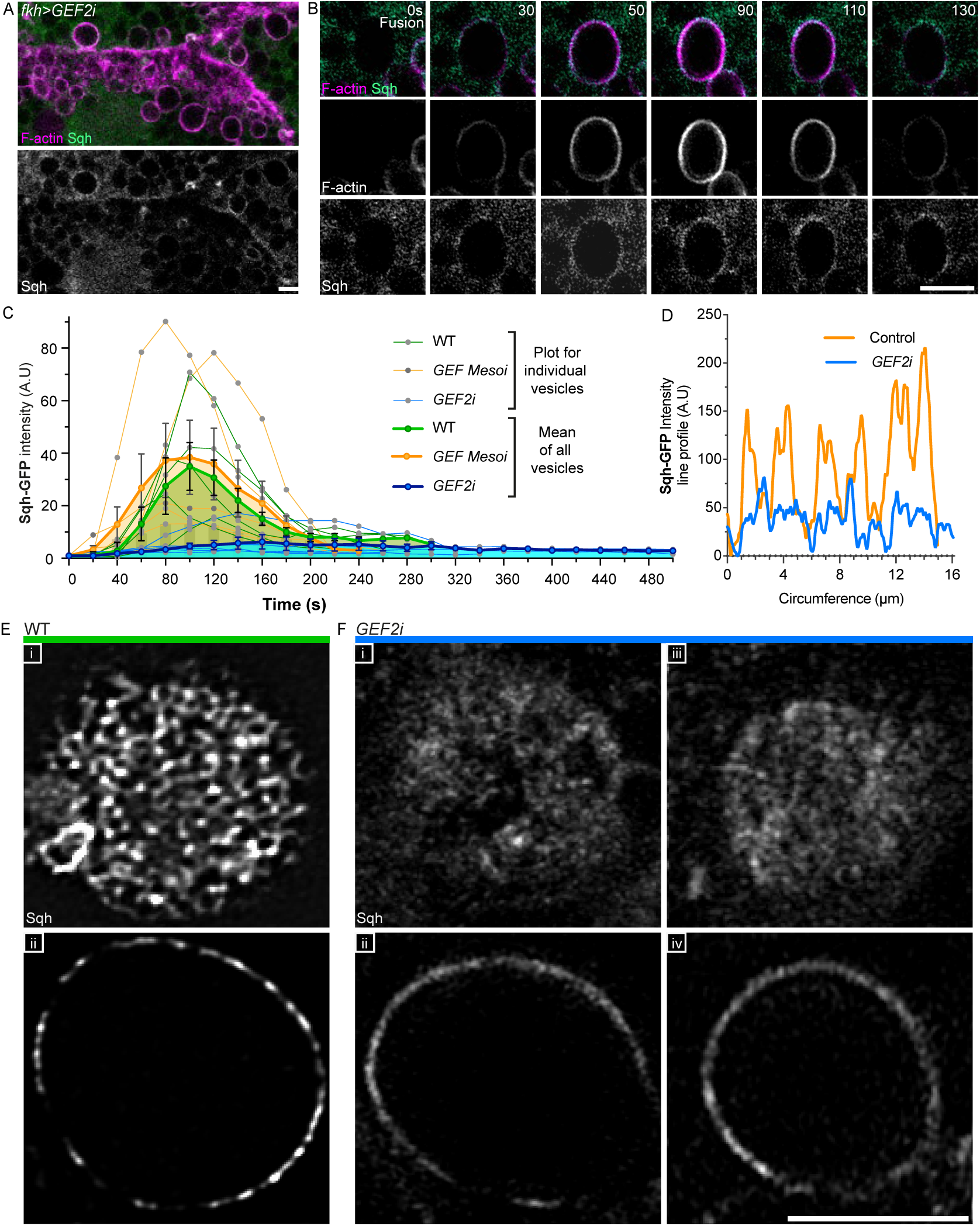
RhoGEF2 is essential for myosin II recruitment and organization on the vesicle. A. Low magnification image of a cultured *RhoGEF2* knockdown (*fkh-Gal4>UAS-RhoGEF2* RNAi) LSG. F-actin (LifeAct-Ruby, magenta-top panel), myosin II (Sqh-GFP, green-top panel or gray-bottom panel). B. Time series of a fused vesicle in a *RhoGEF2* knockdown LSG. F-actin (LifeAct-Ruby, magenta-top panel or gray-middle panel), myosin II (Sqh-GFP, green-top panel or gray-bottom panel). Vesicle constriction was stalled (in 72±12% of cases in this background), and the levels of Sqh-GFP recruited to the vesicles appeared lower than in WT (compare to Fig. 1 C). C. Plot of total Sqh-GFP levels on fused SVs over time. Levels were normalized to that of the first appearance of a signal (typically at 40s). Individual SVs are shown for WT, *RhoGEFmeso* RNAi and *RhoGEF2* RNAi (green, orange and blue thin lines, respectively). The mean intensity profile (n=5 vesicles, from 3 LSGs, error bar SEM) is shown by a green, orange and blue fill, respectively. Sqh-GFP levels were dramatically reduced in *RhoGEF2*-RNAi vesicles. D. Representative myosin II (Sqh-GFP) fluorescence intensity line profiles along the circumference of a vesicle from *fkh>RhoGEF2* RNAi, shown in B at the 50s time point (blue line), and a control vesicle from *fkh>RhoGEFmeso* RNAi shown in Fig. S3 A, at the 60s time point (orange line). Sqh-GFP was recruited at lower levels and in a diffuse fashion in vesicles from *RhoGEF2* knockdown LSGs. The florescence intensity distribution along the vesicle circumference between *RhoGEF2* RNAi and *RhoGEFmeso* RNAi controls was found to be significantly different (Kolmogorov-Smirnov test, p value < 0.0001, Kolmogorov-Smirnov D= 0.9689; n=6 vesicles from 3 LGSs from each group). The variance in the amplitude of intensity along the vesicle circumference is a proxy for myosin patterning and was found to be significantly different between the *RhoGEF2* RNAi and *RhoGEFmeso* RNAi controls (for the vesicles represented here, non-parametric Levene’s test, p value= 2.06238E-40, F value= 201.48, F critical= 3.85). E-F. Organization of myosin II (Sqh-GFP) on a WT vesicle and on two *RhoGEF2* knockdown (*fkh-Gal4>RhoGEF2* RNAi) vesicles following fixation and GFP antibody staining. Top panels represent the 3D organization of myosin II on the vesicle surface while the bottom panels represent the distribution at the equatorial plane. In the surface view, the majority of WT vesicles exhibited a discrete, structured pattern of Sqh-GFP (E i, 87%, n=30 vesicles, from 5 LSGs). However, on *RhoGEF2* knockdown vesicles, Sqh-GFP appeared at reduced levels and in a dispersed manner. Specifically, 30% of vesicles showed partial loss of the Sqh pattern, with intermittent gaps in the dispersed arrangement (F i) and another 50% displayed a fully dispersed Sqh arrangement (F iii, n=30, from 5 LSGs). Consistently, at the equatorial plane, the *RhoGEF2* knockdown vesicles showed a more uniform and diffuse pattern of Sqh (F ii, iv) as compared to the structured pattern in WT (E ii). Scale bars: 5 µm (A-B, E-F).

In conclusion, while compromising the levels of RhoGEF2 did not affect actin polymerization, a marked effect on myosin II recruitment to the vesicles was observed. We propose that fusion of the vesicle with the apical membrane leads to the appearance of initial activated Rho1 at the vesicle membrane by diffusion (Fig. S3 C, D). This initial level of active Rho1 is sufficient for the recruitment of Dia and induction of actin nucleation, but cannot drive proper myosin II recruitment and activation on its own. The subsequent activity of RhoGEF2 is required to promote the recruitment of high levels of myosin II in a meshwork pattern.

### RhoGEF2 spatio-temporal organization dictates the recruitment of myosin II

To clarify the role of RhoGEF2 for appropriate myosin II recruitment, we first determined the spatio-temporal pattern of RhoGEF2 on the SVs. Live imaging showed that RhoGEF2-sfGFP was recruited to the vesicle from the cytoplasm in a punctate pattern, typically between 25s to 30s after fusion (Fig. 4 A). RhoGEF2 recruitment preceded myosin II appearance (monitored by Sqh-RFP) by several seconds, suggesting a possible causal relationship (Fig. 4 A). This notion is further supported by the observation that RhoGEF2 knockdown LSGs displayed only low levels of activated Rho1, lacking the typical WT punctate organization (Fig. 2 D, E). Thus, we conclude that the prominent signal of activated Rho1 in WT vesicles can be attributed to its activation by RhoGEF2.

**Figure 4.**
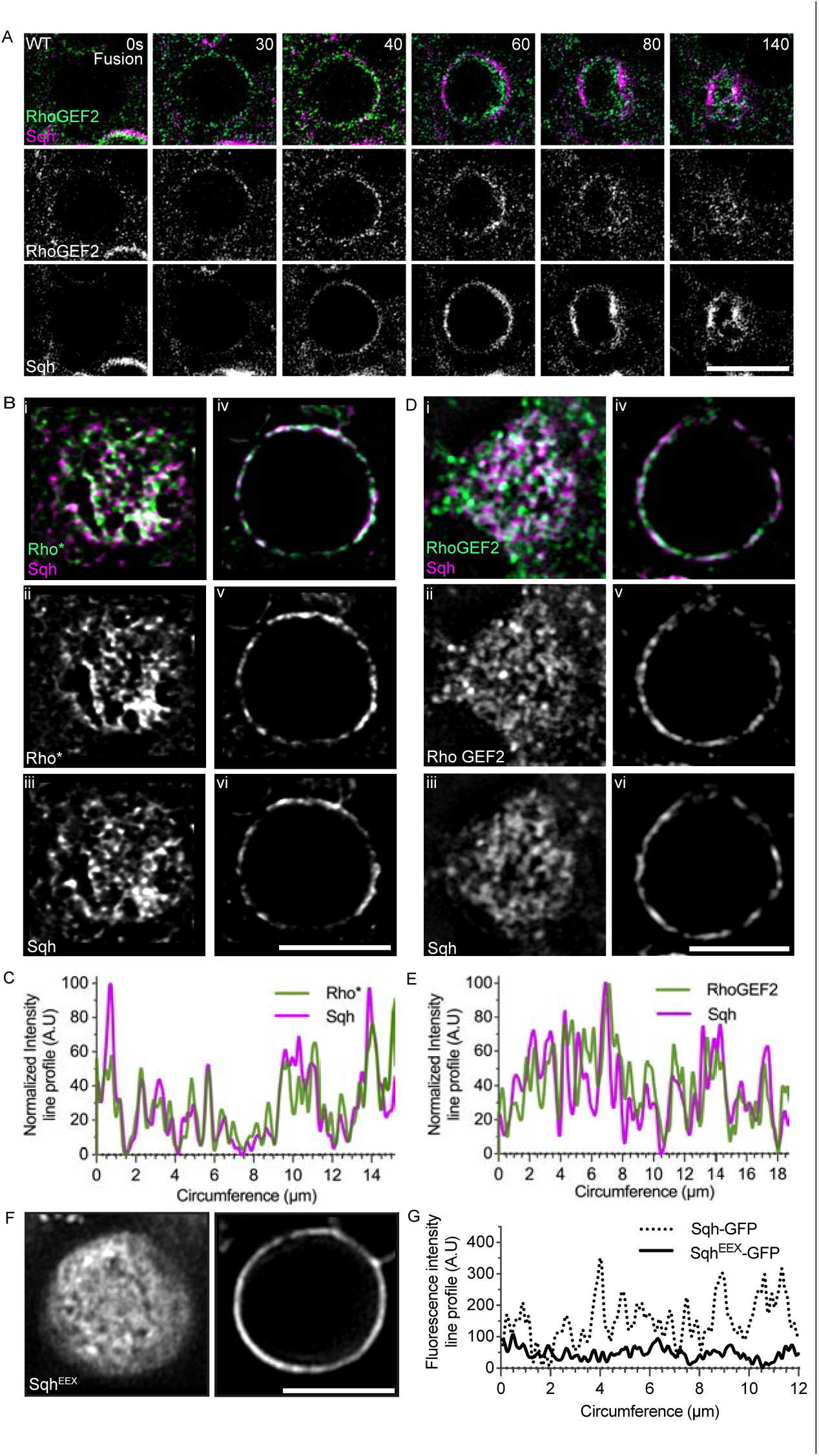
RhoGEF2 spatiotemporal organization directs the recruitment of myosin II. A. Time series of a WT vesicle showing RhoGEF2-sfGFP (green-top panel or gray-middle panel) and Sqh-RFP (magenta-top panel or gray-bottom panel). Recruitment of RhoGEF2-sfGFP was observed at 30s post fusion, while appearance of Sqh-RFP was detected starting at 40s post fusion. Both persisted and displayed a structured appearance throughout the process of vesicle constriction. B. Surface (left, i-iii) and equatorial plane (right, iv-vi) views of a fixed WT vesicle stained for active Rho sensor (green-top panel or gray-middle panel) and Sqh-RFP (magenta-top panel or gray-bottom panel). Both displayed a partially overlapping discrete distribution. C. Fluorescence intensity line profiles from the equatorial plane of the vesicle in B. The intensities of the two reporters were highly correlated. D. Surface (left, i-iii) and equatorial plane (right, iv-vi) views of a fixed WT vesicle stained for RhoGEF2-GFP (green-top panel or gray-middle panel) and Sqh-RFP (magenta or gray). Both displayed a partially overlapping discrete distribution. E. Fluorescence intensity line profiles from the equatorial plane of the vesicle in D. The intensities of the two reporters were highly correlated. F. Surface (left) and equatorial plane (right) views of a fixed WT fused vesicle expressing phosphomimetic myosin, Sqh^EEX^, tagged with GFP and stained with anti-GFP. A fairly uniform recruitment was observed, consistent with constitutive recruitment of Sqh^EEX^-GFP being independent of any underlying molecular asymmetry, including that set up by Rho signaling. G. Plot showing Sqh-GFP (dotted line) and Sqh^EEX^-GFP (solid line) fluorescence intensity profiles along the circumference of representative fused vesicles imaged live. The intensity at each point was normalized to the minimum intensity along the circumference. The variance in the amplitude of intensity of Sqh^EEX^-GFP vs Sqh-GFP, which is measure of the patterning, was found to be significantly different (for the vesicles represented here, non-parametric Levene’s test, p value= 2.6138E-40, F value= 206.86, F critical= 3.86). The pattern of Sqh^EEX^-GFP appeared dispersed in 56+11% of vesicles (n=30, from 3 LSGs). Scale bars: 5 µm (A, B, D, F).

We next monitored the spatial distribution of RhoGEF2 and activated Rho1. The pattern of activated Rho1 showed a similar structured distribution to that of RhoGEF2 (Fig. 4 B (ii, v) vs. D (ii, v)). However, we could not compare them in the same vesicle, since RhoGEF2 and Rho1 are only available as GFP fusion proteins. Because high levels of activated Rho1 are expected to recruit Rock and myosin II, we examined, as an alternative, the correspondence between the distribution of activated Rho1 and myosin II on the same vesicle. Indeed, a significant correlation between these distributions was observed, suggesting a causal relationship between punctate high levels of activated Rho1 and the corresponding recruitment of myosin II (Fig. 4 B, C).

If the structured RhoGEF2 distribution dictates the pattern of myosin II recruitment, we would also expect to see a correspondence between these patterns. By comparing the distribution of RhoGEF2 and myosin II on the same vesicle, we observed a similar distribution and a significant correspondence of the patterns at the equatorial sections (Fig. 4 D, E). These results suggest that local puncta of RhoGEF2 recruitment from the cytoplasm generate distinct foci of activated Rho1, which, in turn, lead to the corresponding structured pattern of myosin II recruitment.

Is the meshwork recruitment of myosin dictated exclusively by the pattern of RhoGEF2, or are there structural spatial asymmetries on the vesicle which may generate a pre-pattern for myosin II recruitment? To examine this possibility, we followed the distribution of a phospho-mimetic myosin protein (Sqh^EEX^), which is constitutively active and does not require phosphorylation by Rock (Munjal *et al*., 2015). The localization of Sqh^EEX^ thus bypasses the normal recruitment signals. Sqh^EEX^ localized in a fairly uniform pattern in the majority of the vesicles (Fig. 4 F, G) and led to partial stalling of vesicle constriction (39±5%, n=75 SVs, N=3 LSGs). This result implies that apart from the local patterning by RhoGEF2, there are no other inherent molecular or physical asymmetries on the SV membrane that impinge on the recruitment of activated myosin II. We conclude that the punctate distribution of RhoGEF2 dictates the spatial pattern of activated Rho and Rock in WT SVs, which subsequently drive the structured pattern of myosin II recruitment.

We next examined what triggers the recruitment of cytoplasmic RhoGEF2 to the vesicles. To determine whether the recruitment of RhoGEF2 to the vesicle depends on the polymerization of F-actin, we followed its recruitment in LSGs treated with the actin polymerization inhibitor Latrunculin A (LatA; Fig. 4S). Fused vesicles were marked and detected with the F-actin reporter LifeAct-Ruby, or a membrane marker CAAX-mCherry. Prior to the addition of LatA, the normal punctate recruitment of RhoGEF2 was observed (Fig. 4S A, B). However, following addition of LatA, newly fused SVs were stalled and displayed an irregular F-actin pattern of a few puncta. No recruitment of RhoGEF2 to the SVs was observed (Fig. 4S C, D). This experiment indicates that F-actin, or an actin-associated protein, is required to recruit RhoGEF2 to the fused SVs. Such a mechanism would ensure the pre-establishment of F-actin tracks for the myosin II motors to assemble.

### The punctate distribution of myosin II drives constriction and crumpling of secreting vesicles

Having established the basis for the punctate recruitment of myosin II, we next wanted to explore its role in vesicle constriction. Glue SVs do not undergo isotropic constriction, but rather demonstrate multiple local folds in a process we have termed “crumpling” (Kamalesh *et al*., 2021). As a result, in this novel mode of secretion, when the vesicle content is effectively expelled, the vesicle membrane remains insulated from the apical cell membrane. Could the focal recruitment of myosin II direct the process of vesicle crumpling?

To examine this notion, we followed the spatial correlation between the distribution of myosin II and the topology of the constricting vesicle membrane. In cross sections of a squeezing vesicle, the sites of high myosin concentration indeed corresponded to the location of membrane folding (Fig. 5 A), suggesting a causal relation between local myosin recruitment and the sites of vesicle membrane folding. The actual squeezing of the vesicle content is a fairly rapid event that occurs within a timeframe of 36±8 s within the vesicle fusion-secretion window (Kamalesh *et al*., 2021); we propose that it is executed by the concerted action of multiple confined actomyosin constriction events, that are dictated by the focal recruitment and distribution of myosin II. Local constrictions of the actomyosin mesh lead to buckling of the underlying vesicle membrane. The combination of multiple local events leads to the global squeezing of the vesicle, content release, and crumpling of the vesicle membrane (Fig. 5 B).

**Figure 5.**
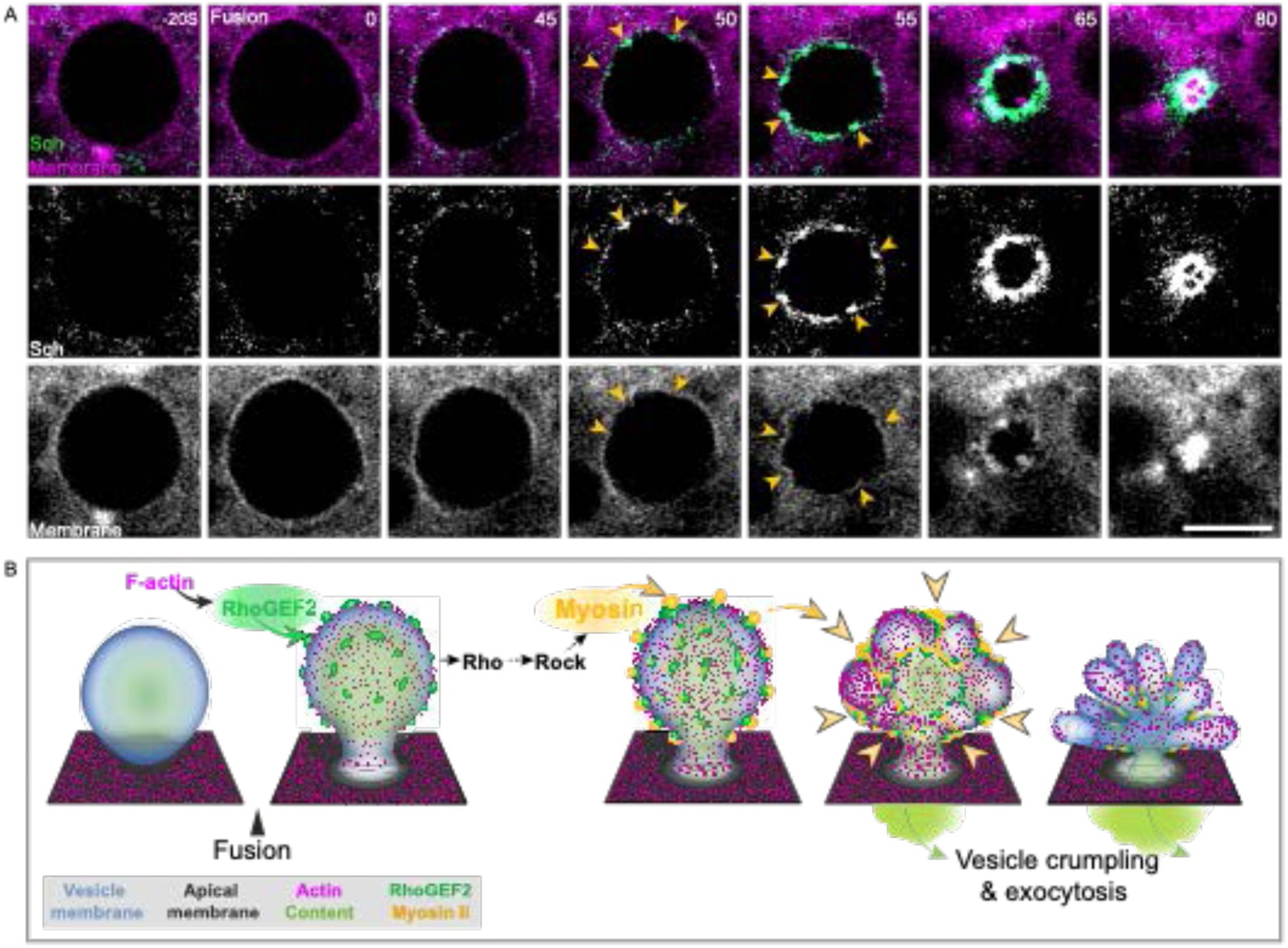
Punctate distribution of myosin II drives constriction and crumpling of secreting vesicles. A. Time series of a constricting WT vesicle, showing Sqh-RFP (green-top panel or gray-middle panel) and the mCD8-GFP membrane marker (magenta-top panel or gray-bottom panel). Folding of the vesicle membrane at several sites was observed. Distinct foci of Sqh-GFP were detected above the prominent membrane folds (arrowheads). This observation suggests that high myosin II levels drive local constriction of actomyosin, leading to vesicle membrane buckling. Scale bar: 5 µm B. Schematic of orchestrated actomyosin assembly on large secretory glue vesicles. Following vesicle fusion with the apical membrane, activated Rho1 diffuses to the vesicle membrane and recruits Dia, which drives uniform F-actin nucleation (magenta). The F-actin coat provides an unknown signal to recruit RhoGEF2 (green) from the cytoplasm, in a structured pattern. RhoGEF2 boosts the level of activated Rho1 at these foci, which in turn, act to recruit myosin II (yellow) via Rock. The enhanced levels of myosin II on the vesicle and its structured distribution drive vesicle constriction at multiple sites, giving rise to both content expulsion and crumpling of the vesicle membrane. Subsequently, clathrin-mediated endocytosis clears the residual SV membrane.

## Discussion

We have monitored the mechanism of actomyosin recruitment to the large glue secretory vesicles in the *Drosophila* larval salivary gland, following their fusion with the apical cell membrane. This process extends over 3-4 minutes for each vesicle, culminating in the complete expulsion of vesicle cargo into the LSG lumen. Contraction of an actomyosin network that coats the vesicle plays an essential role in content release, leading us to explore in depth the molecular mechanisms underlying its formation and activation. In keeping with a standard and well-established pathway leading to actomyosin-based contractility, both F-actin assembly via the Formin Dia and myosin II recruitment by Rock are triggered in this setting by activated Rho1. However, our observations reveal that the spatiotemporal characteristics of the two responses to active Rho1 are distinct.

Prior to fusion, the secretory vesicle membrane is devoid of activated Rho1. Fusion of the vesicle can allow diffusion of membrane-tethered active Rho1 from the apical membrane to the vesicle membrane. Diffusion of active Rho1 to the vesicle membrane is arrested after formation of the actin coat (Kamalesh *et al*., 2021). This initial, low level of active Rho1, uniformly distributed 1 on the vesicle membrane, leads to a corresponding pattern of recruitment of the Formin Dia and the subsequent fairly even formation of prominent actin cables, but for only low levels of myosin II recruitment, which are insufficient to drive vesicle constriction. Accumulation of F-actin on the vesicle, or of hitherto unknown actin-associated proteins, leads to the recruitment of RhoGEF2 from the cytoplasm to the vesicle membrane. This recruitment is uneven and marked by multiple local puncta of high RhoGEF2 levels, which drive local Rho activation and subsequent recruitment of myosin II (mediated by Rock) in a corresponding structured pattern. The distribution and activity of myosin II then drive the constriction of actomyosin at multiple locations, leading to buckling of the underlying vesicle membrane. These constrictions enable the unique secretion mode from such large vesicles (termed exocytosis by vesicle “crumpling”), which expels the vesicle content, while retaining the vesicle membrane insulated from the apical membrane of the LSG (as shown schematically in Fig. 5 B).

Activated Rho has been universally demonstrated to be the critical switch for recruitment of both F-actin and myosin II (Hodge & Ridley, 2016; Jaffe & Hall, 2005; Spiering & Hodgson, 2011), but the temporal aspects of activation of the two responding branches have not been explored in detail. We identified a biphasic recruitment mode in the *Drosophila* LSGs, where the activation of myosin II takes place after the formation of a filamentous actin coat, and may actually rely on components generated by actin coat construction. The temporal basis for the biphasic recruitment in this system stems from an apparent difference in the levels of active Rho1 required to trigger each branch. Low levels of activated Rho1, provided by diffusion from the apical membrane, are sufficient for full-fledged recruitment of Dia and subsequent nucleation and polymerization of an F-actin coat. However, they are insufficient for complete myosin II recruitment, which relies on an enhanced second wave of Rho1 activation, triggered by RhoGEF2 activity. In light of our observations, examining whether the two branches are activated simultaneously or consecutively in other systems of actomyosin recruitment, and if they have distinct Rho activation thresholds is warranted. For example, such a mechanism may trigger the recently identified structured recruitment of myosin II to secretory vesicles of mouse salivary glands (Ebrahim *et al*, 2019).

The biphasic mode of actomyosin recruitment is critical for the proper secretory activity of the salivary gland. Without the second wave of myosin II recruitment, triggered by RhoGEF2, only low levels of myosin II accumulate on the vesicles and they remain stalled. This regulatory mode serves two roles for the constricting vesicles. First, it allows harnessing the high levels of inactive RhoGEF2 in the cytoplasm, by recruiting the protein and activating it at the vesicle membrane, presumably via G-protein-coupled receptors (GPCRs) (van Unen *et al*, 2015). Second, the recruitment and distribution of RhoGEF2 to the vesicles rely on elements that accumulated in the first wave of activation, namely F-actin or actin-associated proteins. These unevenly distributed local foci lead to a corresponding accumulation of RhoGEF2. The structured distribution of subsequent myosin II clusters dictates the constriction pattern of the vesicle, such that the underlying membrane buckles to release the vesicle content but does not integrate into the apical membrane of the cell. This mode of constriction is robust and efficient since dozens of such myosin II foci drive the constriction of each vesicle, while their precise number and spacing can vary between vesicles. The resulting outcome of membrane crumpling maintains the unique composition of the apical membrane in the face of fusion to multiple large secretory vesicles (Kamalesh *et al*., 2021).

## Acknowledgments

We thank Dr. R’ada Massarwa and Vandana Bharadia for help at the initial stages of this work. The research was supported by Israel Science Foundation (grant no. 706/20) to B.-Z.S., O.A., and E.D.S. and the Minerva Foundation with funding from the Federal German Ministry for Education and Research. O.A. also acknowledges funding from the Henry Chanoch Krenter Institute for Biomedical Imaging and Genomics, the Schwartz Reisman Collaborative Science Program, the Yeda-Sela Center for Basic Research, and the European Research Council (ERC) under the European Union’s Horizon 2020 research and innovation program (grant agreement no 851080). O.A. is an incumbent of the Miriam Berman presidential development chair. B.-Z.S. is an incumbent of the Hilda and Cecil Lewis Professorial Chair in Molecular Genetics.

This paper is dedicated to the memory of the late Dr. R’ada Massarwa, a groundbreaking scientist with an unlimited passion to visualize and uncover the wonders of Biology.

## Author contributions

B-Z.S., O.A., E.D.S. and K.K. conceived and designed the experiments. K.K. carried out most of the experiments and analyzed the data, D.S. performed the initial experiments and analyzed the data. B-Z.S., O.A., E.D.S. and K.K. wrote the manuscript.

---

**Figure S1.**
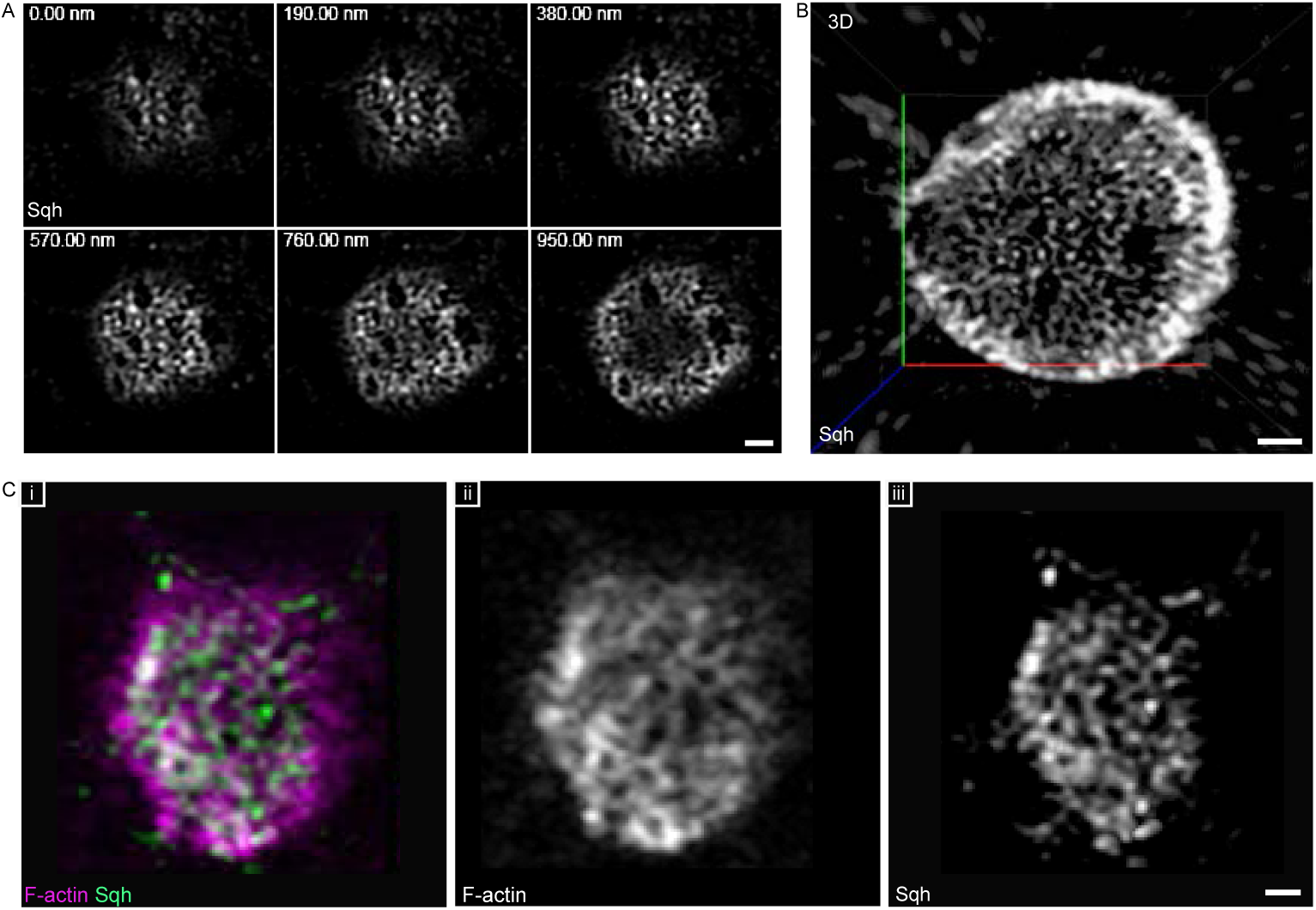
3D organization of actomyosin on an LSG secretory vesicle. A. Airyscan confocal images of the Z-planes used to construct the surface view of Sqh-GFP staining shown in Fig. 1 G. B. 3D reconstruction of the Sqh-GFP distribution on the entire vesicle, as viewed through the vesicle fusion pore. C. Surface view showing F-actin (phalloidin) staining (magenta- (i) or gray- (ii)) and Sqh-GFP staining (green- (i) or gray- (iii)). F-actin is spread-out over the entire surface of the vesicle in a non-homogenous manner, while Sqh is distinctly patterned as a discrete meshwork. Scale bars: 1 µm (A-C).

**Figure S2.**
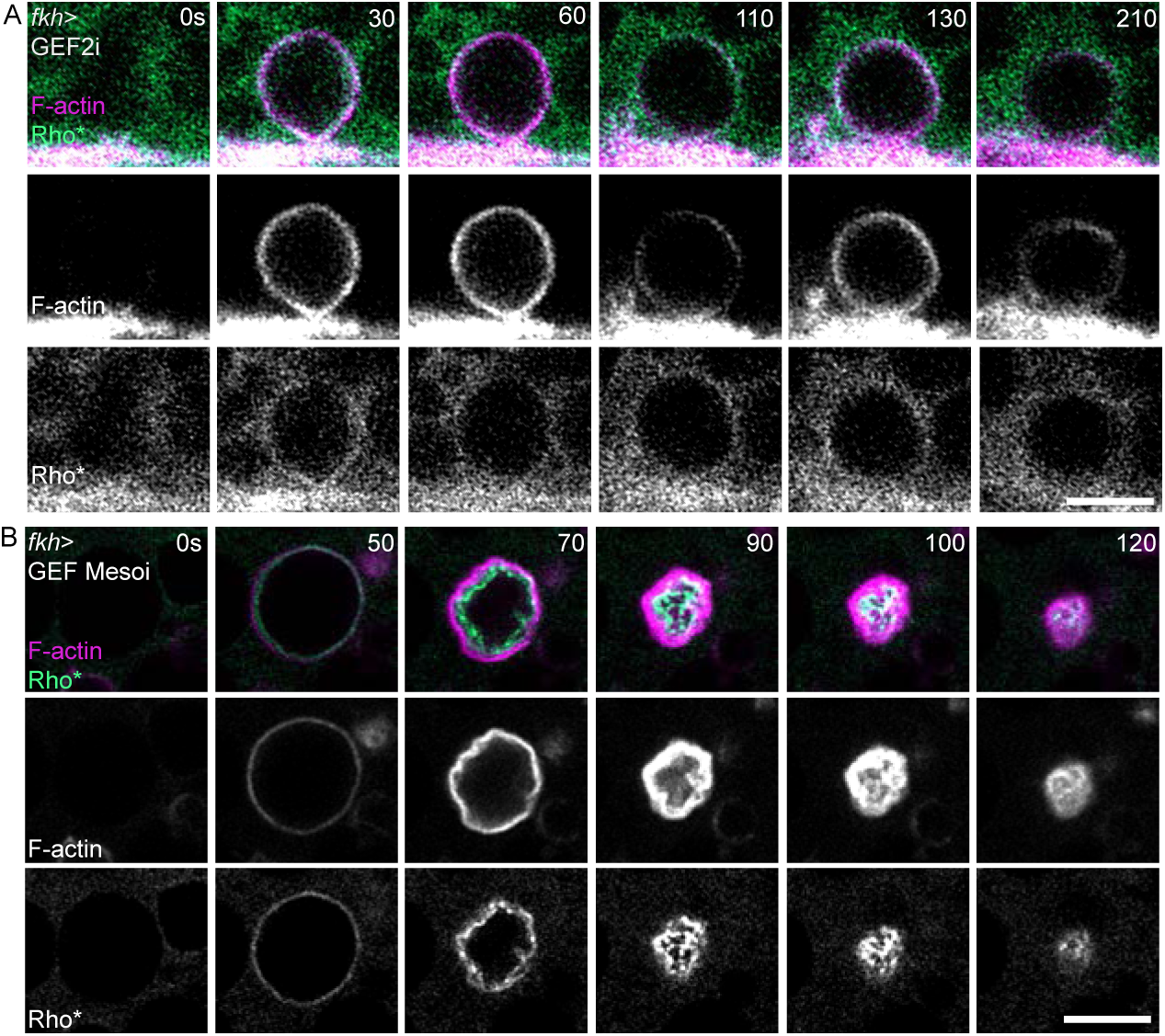
Knockdown of *RhoGEF2* affects activated Rho levels on LSGs. A. Time series of a representative vesicle from a *RhoGEF2* knockdown LSG (*fkh-Gal4>UAS-RhoGEF2* RNAi), displaying the typical F-actin assembly and disassembly cycle (monitored using Lifeact-Ruby, magenta-top panel, or gray-middle panel), similar to WT contraction-halted vesicles (treated with a Rock inhibitor). This vesicle failed to constrict, and the levels of recruited active Rho sensor (Rho*, green-top panel or gray-bottom panel) were low and diffuse (compare to WT, Figure 2C). Of all the stalled vesicles in *RhoGEF2* RNAi-expressing glands, 22% exhibited the typical F-actin cycling behavior. B. Time series of a representative vesicle from a *GEFmeso* knockdown LSG (*fkh-Gal4>UAS-GEFmeso* RNAi), displayed normal constriction dynamics, and active Rho1 sensor (Rho*, green-top panel or gray-bottom panel) and F-actin (Lifeact-Ruby, magenta-top panel or gray-middle panel) recruitment features, similar to those observed in WT. Scale bars: 5 µm (A-B).

**Figure S3.**
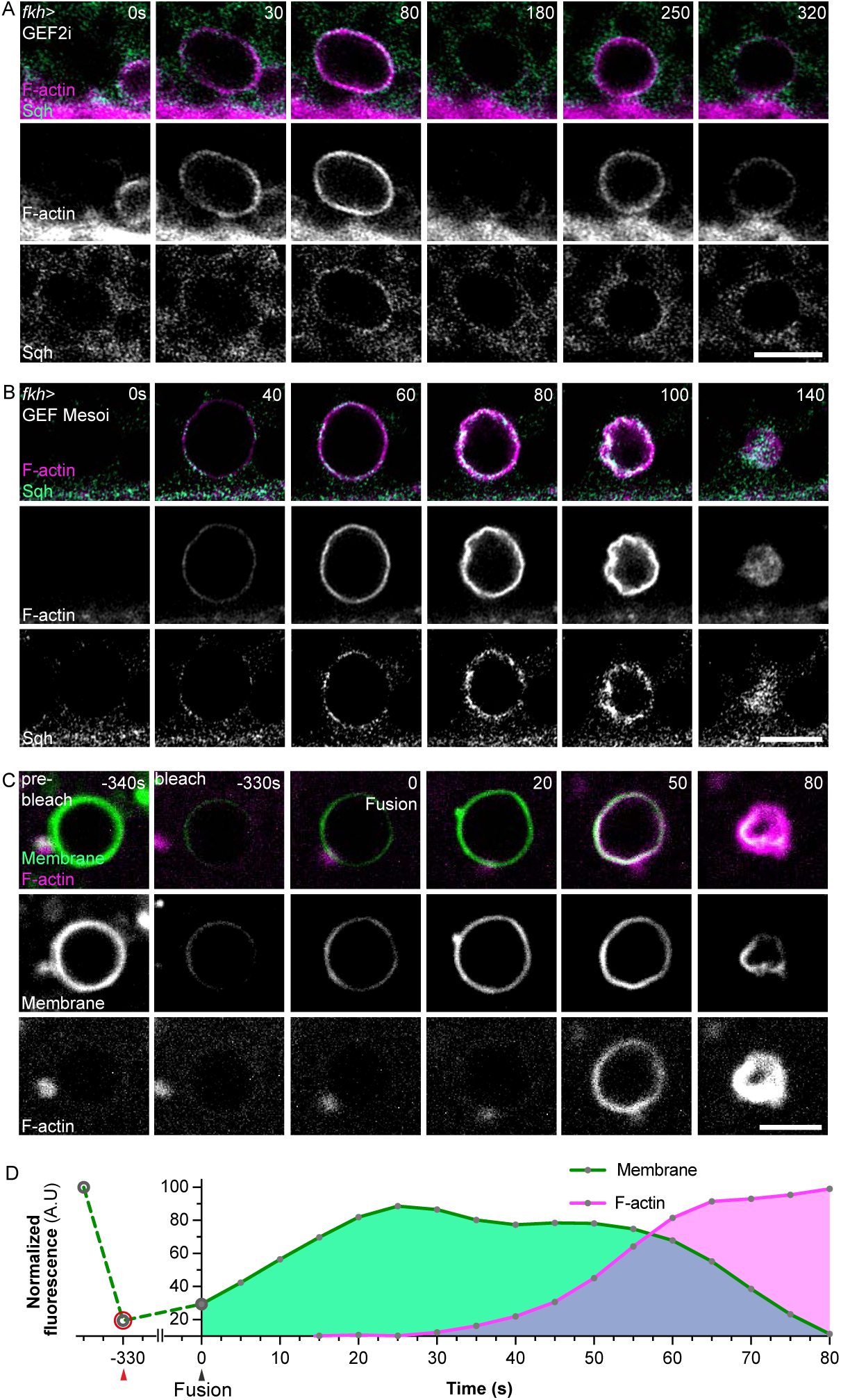
RhoGEF2 affects myosin II recruitment. A. Time series of a representative vesicle from *RhoGEF2* knockdown LSG displaying the typical F-actin assembly and disassembly cycle (monitored using LifeAct-Ruby, magenta-top panel or gray-middle panel), similar to that observed in WT Rock inhibitor-treated contraction-halted vesicles. This vesicle failed to constrict and the levels of recruited myosin II (Sqh-GFP green-top panel or gray-bottom panel) were low and diffuse (compare to WT, Fig. 1 C). B. Time series of a representative vesicle from *GEFmeso* knockdown LSG, showing the typical F-actin (LifeAct-Ruby, magenta-top panel or gray-middle panel) and myosin II (Sqh-GFP, green-top panel or gray-bottom panel) recruitment features similar to WT. C. Transient diffusion of membrane-associated molecules following vesicle fusion. Time-series showing FRAP of a membrane reporter (mCD8-GFP, green-top panel or gray-middle panel) at vesicle fusion. Photobleaching of mCD8-GFP was performed on the vesicle before fusion (-330s) and monitored over time. F-actin (LifeAct-Ruby, magenta-top panel or gray-bottom panel) assembled after vesicle fusion and remained during vesicle constriction. At the instance of vesicle fusion (identified by expansion of the vesicle size) but prior to appearance of F-actin, mCD8-GFP fluorescence recovery was observed, indicating a transient time interval during which diffusion occurs. D. Fluorescence recovery profile for mCD8-GFP at the instance of vesicle fusion, for the time-series shown in C. Photobleaching is denoted by the red arrowhead and vesicle fusion by the black arrowhead. The fluorescence intensity at each point was normalized to the first appearance of the signal. Scale bars: 5 µm (A-C).

**Figure S4.**
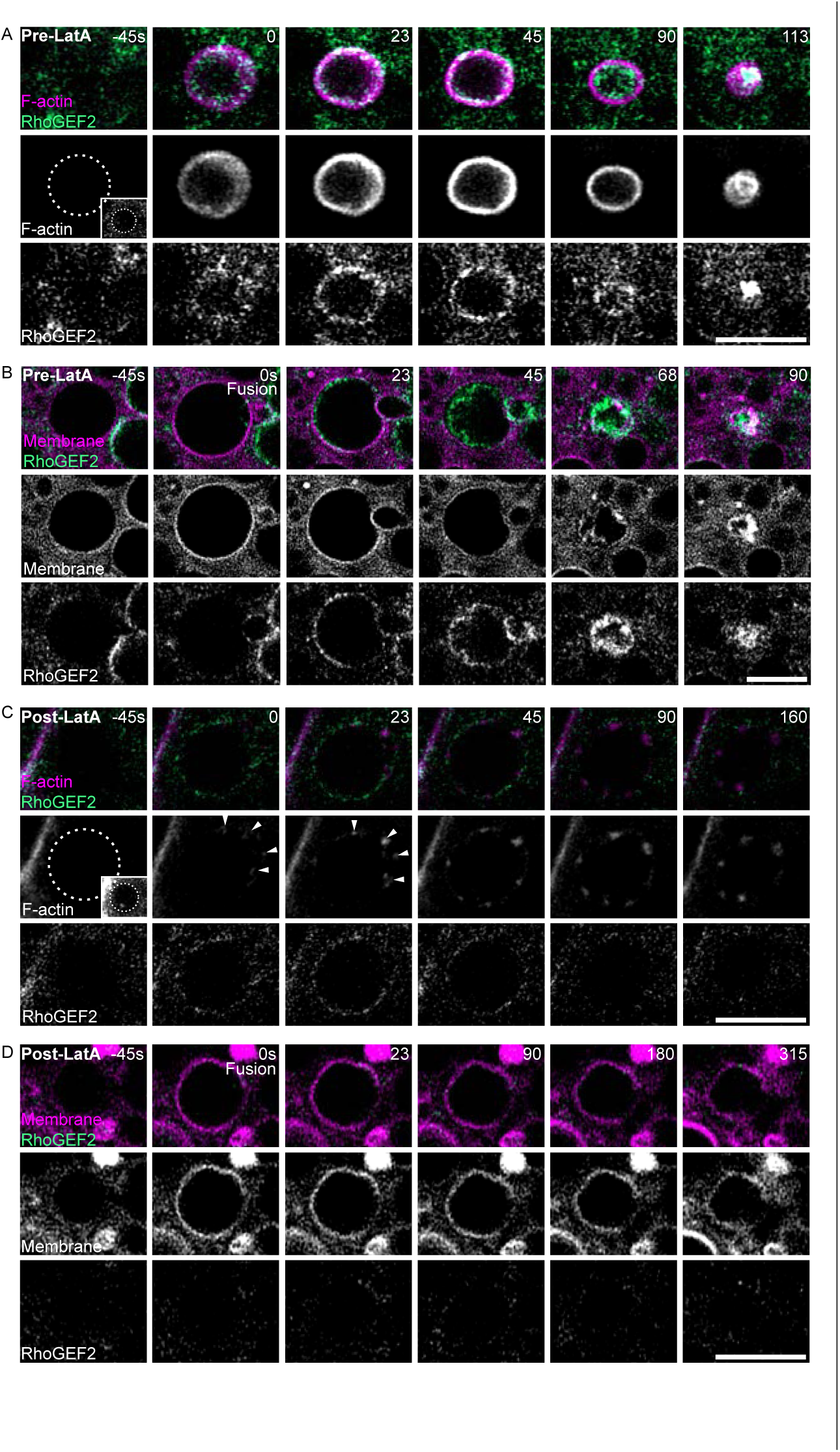
GEF2 recruitment to the fused secretory vesicle is dependent on F-actin polymerization. A. Time series of a representative WT secreting vesicle showing the temporal recruitment of RhoGEF2 (RhoGEF2-sfGFP, green-top panel or gray-bottom panel) with respect to F-actin coat assembly (LifeAct-Ruby, magenta-top or gray-middle), following vesicle fusion. Time 0s corresponds to when F-actin assembly becomes clearly visible, previously characterized as ∼40s post-vesicle fusion (Kamalesh *et al*., 2021). The dotted line in the inset in the middle plane represents the approximate outline of the vesicle at the time of fusion (deduced from expansion in vesicle size). B. Time series of a representative WT secreting vesicle showing the recruitment dynamics of RhoGEF2 (GEF2-sfGFP green-top panel, gray-bottom panel) when following the vesicle using the membrane marker CAAX-mCherry (magenta-top panel, gray-middle panel). The transient expansion of the vesicle (compare vesicle diameter at 0s with -45s) and the appearance of membrane marker on the vesicle (likely due to its diffusion from the apical surface), indicates the onset of vesicle fusion that corresponds to time 0s. C. Time series of a representative secretory vesicle from the same LSG as the vesicle shown in (A), post treatment with latrunculin-A (LatA), an inhibitor of F-actin polymerization. Adding LatA to the medium prevented F-actin coat assembly on the vesicle and RhoGEF2 recruitment to vesicles. Fused vesicles were identified by the appearance of a few F-actin clusters post-fusion (pinpointed by arrowheads in the middle panel). The dotted line in the inset at the middle plane represents the approximate outline of the vesicle at the time of fusion (which can be observed when the fluorescence is scaled appropriately). D. Time series of a representative secreting vesicle from the same LSG as the vesicle shown in (B), post-treatment with LatA. The vesicle was visualized using the membrane marker CAAX-mCherry that highlights it post-fusion. Addition of LatA prevented RhoGEF2 recruitment to the vesicles. Scale bars: 5 µm (A-D)

## Materials and Methods

### *Drosophila* strains and rearing conditions

*Drosophila* fly lines used in this study include: Lines obtained from the Bloomington Drosophila Stock Center (NIH P40OD018537): *fkh*-GAL4(B-78060; Figs. 1-5 & Extended Data Figs. 1-5), UAS-LifeAct-Ruby (B-35545; Figs. 1-5 & Extended Data Figs. 1-5), *sqh*–GFP.RLC (B-57145; Fig. 1, 3, 5 & Fig. S1-S5), UAS-*RhoGEF2* RNAi (B-34643; Figs. 2, 3 & Fig. S2, S5 and B-31239 data not shown), UAS-*GEFmeso* RNAi (B-42545; Figs. 2, 3 & Fig. S2, S3), *sqh*-mCherry (B-59024; Fig. 4 & Figs. S2, S5), UAS-mCherry-CAAX (B-59021; Fig. 5), *sqh*-sfGFP-*RhoGEF2* (B-76260, Fig. 4). Lines from other resources: *ubi*-AniRBD::GFP (Rho1 sensor; (Munjal *et al*., 2015) kindly provided by Thomas Lecuit, IBDM (Figs. 2,4); *sqh*E20E2–GFP (phosphomimetic Sqh (Royou *et al*, 2002); kindly provided by Andrea Brand, Univ. of Cambridge, Fig. 4).

All fly stocks were reared on standard cornmeal, molasses and yeast media at 21°C in a temperature-controlled room. Crosses and flies used for imaging experiments were grown in 25°C incubators without internal illumination. Live imaging experiments were performed on *ex-vivo* cultures of third-instar *Drosophila* LSGs. Larvae from crosses were used without distinguishing between sexes, as no obvious sex-specific differences in SG secretion were observed.

### Culturing third instar SGs for live imaging

SG culturing was performed as previously described (Kamalesh *et al*., 2021). In brief, SGs from third instar larvae were dissected out in Schneider’s *Drosophila* medium (SDM) and up to 6 glands were transferred to a 35-mm dish, with a 10 mm #1.5 glass bottom well (Cellvis D35-14-1.4-N) containing 200 μl of fresh medium for live imaging. LSGs were typically imaged within an hour of dissection.

### Drug treatment of LSGs

LSGs were imaged for at least 5 min before drug treatment, to ensure overall health and the commencement of secretion. With image acquisition stopped, LatA (1 µM; Sigma-Aldrich), was added to the medium with a micropipette, directed near but without touching the sample, and mixed gently with a pipette tip. Typically, the effect of LatA was clearly observable after 15mins of its addition. Image acquisition was recommenced on the same region of the LSG to directly compare the state of F-actin (visualized using the Lifeact-Ruby reporter) before and after LatA treatment. Cultures maintained for up to 2hr after treatment did not display any overt abnormalities in tissue integrity.

### Airyscan time-lapse imaging and image processing

Live imaging of the LSGs was performed on an LSM900 Airyscan 2 confocal microscope (Zeiss), using a 63X/1.4 NA (oil) objective, 1.5–2X digital zoom, in super-resolution mode. The high-sensitivity Airyscan GaAsP-PMT detector was used, as it significantly improved the signal-to-noise ratio, and thus spatial and temporal resolution. Special care was taken to monitor the alignment of the Airyscan detector throughout the live imaging session, as the movements inside the live specimen could be a source of fluctuations in the detector alignment. The Airyscan detector adjustments were set to activate alignments automictically during live and continuous scans, and during time-series acquisitions. Time series were acquired from a region close to the lumen of the LSG, where the vesicles fuse. To achieve the highest temporal resolutions, image acquisitions were made from single optical sections with no time intervals and for short durations of 5-10mins only, due to profuse bleaching. For quantifying vesicular dynamics (see section below), two optical sections 2 μm apart were typically acquired with a 5s time interval for a duration of 20mins. For all live Airyscan imaging, scan speeds of ∼ 1µs/pixel were optimal to maintain proper detector alignment while achieving the best temporal and spatial resolution.

Raw stacks were processed using the Airyscan processing utility on Zen 3.1 (blue edition) to obtain the Airyscan time series. Fiji and Adobe Photoshop CC 2021 were used for cropping and adjustment of brightness/contrast of the Airyscan images for visualization purposes. The time series shown in the figures represents the typical dynamics observed in vesicles across more than 6 LSGs.

### Quantification of vesicle dynamics

From the Airyscan time series acquisitions, vesicles that could be clearly followed from the onset of fusion (which is associated with vesicular expansion) were cropped out for analysis. The entire time-lapse focus series was background subtracted using Fiji, before cropping out the individual vesicles for analysis. A region of interest (ROI) was drawn around the perimeter of the vesicle for the probe being analyzed, on each time frame, starting from a frame before fusion and until the end of secretion, or as long as possible in the case of contraction-halted vesicles observed in *fkh*>*RhoGEF2*i LSGs. An auto-thresholding was then applied to the individual channels of the cropped-out vesicles. For analysis of the LifeAct Ruby probe for F-actin, the Isodata type of thresholding was used, while for analysis of the Rho sensor and Sqh intensity, the mean type of thresholding was used. The integrated intensity of the probes post-thresholding and within the drawn ROIs were measured. The measured intensity of each time frame was normalized to the intensity of the time frame when the probe first appeared and plotted over time (Figs. 2 E and 3 C).

### Immunostaining of LSGs

Individual third instar LSGs were dissected out in SDM, then transferred to another drop of SDM supplemented with 1% of freshly prepared PLPS fixative (4% Paraformaldehyde, 0.1% Glutaraldehyde, 0.01M Sodium meta-periodate, 0.075 L-Lysine, 0.035 phosphate buffer, 0.1% Saponin, pH 7.4), where they were swiftly bisected along their length using a pair of fine tweezers, thus exposing the lumen of the glands. The bisected LSGs were immediately transferred to ice-cold PLPS fixative and kept on a rotating shaker for 30 mins at RT. The bisected and fixed LSGs were then given a quick wash with Permeabilization neutralization (PN) solution (0.035 phosphate buffer, 0.01M Sodium meta-periodate, 0.075 L-Lysine, 0.35% Triton-X 100, 0.2% Sodium deoxycholate, 0.2% Glycine, pH 7.4), followed by another 15 min wash on a rotating shaker. Fixed LSGs were pooled together in PLPS buffer (0.035 phosphate buffer, 0.01M Sodium meta-periodate, 0.075 L-Lysine, pH 7.4) until enough glands were obtained, and permeabilized with PN solution for 30 mins at RT followed by blocking in PN solution with 5% normal goat serum (NGS), for 1hr at RT. Primary antibody incubation was performed for 16-24 hrs at 4°C in PBST (PBS+0.3% Triton X-100) with 5% NGS, followed by 3 washes with PBST for 10mins each. Secondary antibody incubation was performed for ∼6 hr at RT or 12-16 hr at 4°C, followed by 3 washes in PBST. Following an additional wash in PBS, the tissue was transferred to 90% glycerol (in PBS) for a couple of hours. The LSGs were then mounted between 2 glass coverslips with spacers, using 90% glycerol as the mounting medium for Airyscan imaging.

The following primary antibodies were used: rabbit anti-GFP (abcam Cat# ab290) diluted 1:200 was used throughout, except when co-staining for mCherry, where chick anti-GFP (abcam Cat# ab13970) was used at 1:800; rat anti-RFP 1:1200 (ChromeTek Cat# 5F8): rabbit anti-dsRed (Clonetech Cat#632496). Anti-rabbit, anti-chick and anti-rat secondary antibodies conjugated to Alexa-488, Alexa-568, or Alexa-633 were purchased from Molecular Probes (Invitrogen) and used as recommended by the manufacturer. Phalloidin-Atto647 (Fluka Cat#65906) or Phalloidin-TRITC (Sigma Cat# P-1951) were used along with primary and secondary antibody incubation when required.

### Airyscan super-resolution imaging for fixed LSGs and image processing

LSGs with their lumens facing close to the coverslip were imaged on the LSM900 Airyscan 2 confocal microscope (Zeiss), using a 63X/1.4 NA (oil) objective, 2-4X digital zoom, in super-resolution mode. Scan speeds >2µs/pixel were optimal for the fixed samples. Around 10-12 μm z stacks were typically acquired using a 0.15 μm interval between optical sections. Multiple channels were acquired in sequential frame acquisition mode. The raw z-stacks were processed using the 3D Airyscan processing utility of Zen 3.1 (blue edition) to obtain Airyscan super-resolution stacks. Fiji and Adobe Photoshop CC 2021 were used for cropping and adjustment of brightness/contrast of the Airyscan images for visualization purposes.

### Analysis of the molecular organization of vesicles

For 3D reconstruction and analysis of vesicles, all processing was done using the Zen 2 (blue edition) service pack. Full Z-stacks of individual vesicles were cropped out from Airyscan Z-stacks of a portion of the gland using the Create image subset utility. The images were further sharpened using the unsharp mask utility, followed by applying a Nearest Neighbor (NN) deconvolution algorithm (adjustable). The contrast was further improved by applying the Enhance contour utility. The 3D reconstruction of individual vesicles obtained with Zen 2 was used to analyze the patterns of myosin II, F-actin, Rho and RhoGEF2 and their overlap in co-immunostaining experiments. Staining of the first 6 sections along the X-Y imaging axis from the surface of the vesicles (irrespective of the orientation of the vesicle) was sum projected, to depict the surface view defined in Fig 1 E. The surface view of myosin II staining in WT is shown in Fig 1. G and in *fkh*>RhoGEF2i in Fig 3 F (i) and G (i). Similarly, surface views of active Rho sensor and RhoGEF2 staining are shown in Fig 4 B (ii) and D (ii), respectively. The surface view gives the impression of the overall 3D pattern of the molecules on the surface of the vesicles.

The equatorial plane view of the vesicles was used to analyze and obtain a quantitative measure of the distribution of myosin II, active Rho sensor and RhoGEF2 and their colocalization. The equatorial plane is the middle section of the vesicle from its Z-stack that is defined in Fig 1 E. The equatorial plane view for myosin II staining on a WT vesicle shows a punctate distribution as in Fig 1 G and Fig 3 E (ii) and in *fkh*>RhoGEF2i a more uniform distribution as in Fig 3 F (i) and G (i). To obtain a quantitative measure of this distribution, line scans depicting the fluorescence intensity along the circumference of the vesicle using the equatorial plane view were plotted using Fiji. For this analysis, the background signal (measured from the center of the vesicle outline) was subtracted from the equatorial plane view, a segmented line of 6-pixel width was drawn along the circumference of the vesicle that encompasses most of the fluorescence signal, and the Fiji plot profile function was used to plot the intensity of the molecules along the vesicle circumference. An example of the line scan plot showing the comparative distribution of F-actin and myosin II in WT is shown in Fig. 1 D. This plot shows large fluctuations in fluorescence intensity distribution for myosin II, characteristic of a punctate or patterned distribution, while F-actin intensity is more uniform along the circumference.

To analyze the overlap between active Rho sensor or RhoGEF2 and myosin II, a line scan depicting the fluorescence intensity of both channels along the circumference of the vesicle from the equatorial plane view was used.

### Quantification and statistical analysis

All experiments for each genetic background and drug treatment were repeated at least three times with glands from different organisms, and representative images/videos are shown. The statistical tests used for each experiment and P values are mentioned in the figure legends. The GraphPad Prism software was used for all statistical analysis and plotting.

